# Control of Spreading Depression with Electrical Fields

**DOI:** 10.1101/234005

**Authors:** Andrew J. Whalen, Ying Xiao, Herve Kadji, Markus Dahlem, Bruce J. Gluckman, Steven J. Schiff

## Abstract

Spreading depression or depolarization is a large-scale pathological brain phenomenon related to migraine, stroke, hemorrhage and traumatic brain injury. Once initiated, spreading depression propagates across gray matter extruding potassium and other active molecules, collapsing the resting membrane electro-chemical gradient of cells leading to spike inactivation and cellular swelling, and propagates independently of synaptic transmission. We demonstrate the modulation, suppression and prevention of spreading depression utilizing applied transcortical DC electric fields in brain slices, measured with intrinsic optical imaging and potassium dye epifluorescence. We experimentally observe a surface-positive electric field induced forcing of spreading depression propagation to locations in cortex deeper than the unmodulated propagation path, whereby further propagation is confined and arrested even after field termination. The opposite surface-negative electric field polarity produces an increase in propagation velocity and a confinement of the wave to more superficial layers of cortex than the unmodulated propagation path. These field polarities are of opposite sign to the polarity that blocks neuronal spiking and seizures, and are consistent with biophysical models of spreading depression. The results demonstrate the potential feasibility of electrical control and prevention of spreading depression.

## MAIN TEXT

### Introduction

Electrical modulation of neural activity with externally applied electric fields (*1–8*) has helped probe and understand brain dynamics, and alter the pathological activity associated with neurological disorders from Parkinson’s disease (*9, 10*) to seizures (*11–13*).

Since the experiments and observations of Rushton (*1*) in single motor neuron axons, externally applied low frequency electric fields have been shown to modulate neuronal excitability. The physics of this modulation has been well studied (*14*), and involves an axis of polarization that exists between the soma and apical dendrites which is affected by external electric fields aligned with the cell geometry (*7*). Electric fields parallel to the somatodendritic axis can depolarize or hyperpolarize the soma (*3*), which leads to changes in the action potential initiation threshold near the soma and the opposite polarization in distal dendrites (*7, 15*). Neurons are susceptible to weak (<1 mV/mm) electric fields (*16*) and neuronal networks have higher sensitivity than single neurons (as low as 140 μV/mm (*17*)). Applied electric fields with 1-100 mV/mm strength have been shown to transiently suppress or excite epileptiform activity in rat hippocampal slices (*11, 12*), speed up, slow down and block cortical traveling waves (*18*), and even modulate epileptiform activity in vivo (*19*).

First observed by Aristides Leão in the cortex of anesthetized rabbits (*20*), spreading depression or depolarization (SD) is a large-scale pathological brain phenomenon related to migraine, stroke, hemorrhage and traumatic brain injury (*21, 22*). It manifests as a slow (2-5 mm/min) traveling wavefront of neuronal depolarization. The wavefront can trigger transient seizures as it encounters fresh brain tissue, and leaves in its wake transiently inactivated and swollen brain. The underlying dynamics of SD are generally regarded as the physiological mechanism of the initial aura in human migraines (*21*). Recent work has demonstrated that there is a unification possible in the modeling of the biophysics of spikes, seizures, and SD, and that each of these dynamics occur within a different state of the neuronal membrane (*23*). Importantly, computational modeling suggests that the ignition of SD is thought to be linked to currents flowing inward through the apical dendrites of neurons (*24, 25*), as opposed to the soma during action potential spike generation.

In pioneering early work, Grafstein found that polarization of the cortical tissue parallel to the direction of a propagating SD wave increased the conduction velocity of SD when the electric field direction was positive to negative in the direction of propagation (*26*). This suggested that the propagation is mediated by the movement of positively charged ions such as potassium. While surface repetitive pulse stimulation has been shown to interfere with the propagation of SD (*26, 27*), the effects of transcortical DC fields on SD has not been well explored.

We note that in the computational and experimental literature on SD, that there appears to be a possible effect of polarizing the neurons along their soma-dendritic axis (*24*), and that the ignition of SD appears to be in the apical dendritic fields rather than at the axon initial segment seen in spike generation (*25*). These findings suggested that polarization of neurons using electrical fields might be able to modulate the ignition or propagation of SD.

In this article, we demonstrate the effects of transcortical DC polarization on SD propagation. We explore the effects of varying field strengths, stimulus timing and methods of SD generation in conjunction with simultaneous intrinsic optical imaging and potassium dye epi-fluorescence. Lastly, we hypothesize plausible mechanisms for the interactions shown and comment on their utility in a clinical setting.

### Results

#### Electric field effects on SD

In order to address our conjecture that polarization along the somatodendritic axis of neurons might modulate SD, we prepared and placed our brain slices with the somatodendritic axis of principal cells (see Figures 1A and 1C) oriented in the direction of the polarizing field. This generates a gradient of charge along the neuronal body from hyperpolarizing at one end to depolarizing at the opposite end. Simultaneous IOS and APG-2 imaging (see Figures 1D-1F) demonstrates the coincidence of the SD intrinsic signal with a concomitant transient decreasing intracellular K+ signal consistent with experimental measurements (*28*) and computational simulations (*23*). Coronal slices maintain the integrity of the full depth of the gray matter, which permits anatomical exploration of polarization effects on cortex in analogous fashion to in vivo electric fields applied to the pial surface where the neuronal somatodendritic axis is aligned with the applied field. For the purposes of this experiment we defined a positive electric field as one which hyperpolarizes the apical dendrites as depicted in Figure 1C (far right), while a negative electric field depolarizes the apical dendrites in the opposite orientation to the depiction in Figure 1C.

**Fig. 1.**
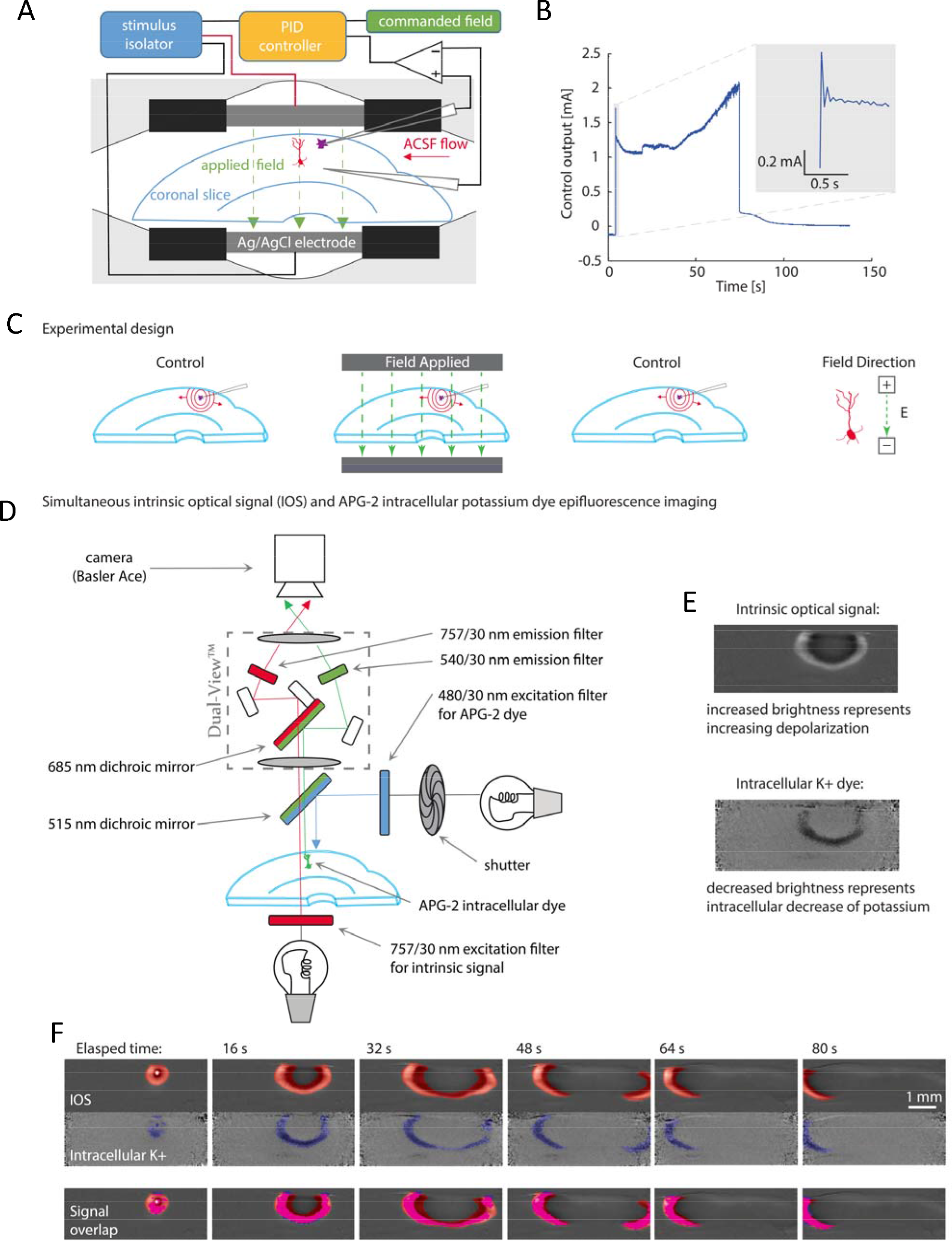
Experimental methods. (**A**) Schematic of the feedback-controlled electric field modified perfusion chamber and coronal slice somatodendtiric axis alignment with the applied transcortical field. Local high K+ is injected from the top pipette (purple) to initiate SD. (**B**) Example control output of the PID feedback field controller, commanding a 116 mV/mm field in the tissue. Inset depicts the slight overshoot performance upon field step on. (**C**) Schematic of a series of 3-epoch experiments consisting of control, field applied, followed by repeat control runs, so that the effect of recovery and time following induction of multiple SDs and the effect of stimulation are accounted for. Also in (**C**) is shown the orientation of the applied surface-positive field with respect to the somatodendritic axis of principal cells in cortex. In (**D**) is shown a schematic of the simultaneous intrinsic optical signal and potassium epifluorescence imaging configuration, utilizing a Dual View multi-channel imaging system (Photometrics, Tucson, AZ), and an example output in (**E**). Simultaneous IOS (red) and APG-2 intracellular potassium dye epi-fluorescence (blue) signals are shown individually, and with overlap (bright pink overlay in (**F**)) indicating that the intracellular K+ decrease during the SD wave is along the leading edge of the IOS signal.

Figure 2 presents the effects of applied DC electric fields on SD induced by local injections of high K+ in the intact cortical gray matter of coronal slices. In Figure 2A, the left column shows a control (unstimulated) trial with SD propagating along all layers of the cortex. The IOS wave in the deeper layers lags behind the propagating wavefront in the superficial layers. The middle column shows that SD propagation is confined to the superficial layers of cortex in the presence of a surface-negative field. In the right hand column, the surface-positive field drives SD propagation towards the deeper layers of cortex where further horizontal spread is arrested and the SD slowly dissipates. See the supplementary data for additional video examples of DC field effects on SD (online only).

**Fig. 2.**
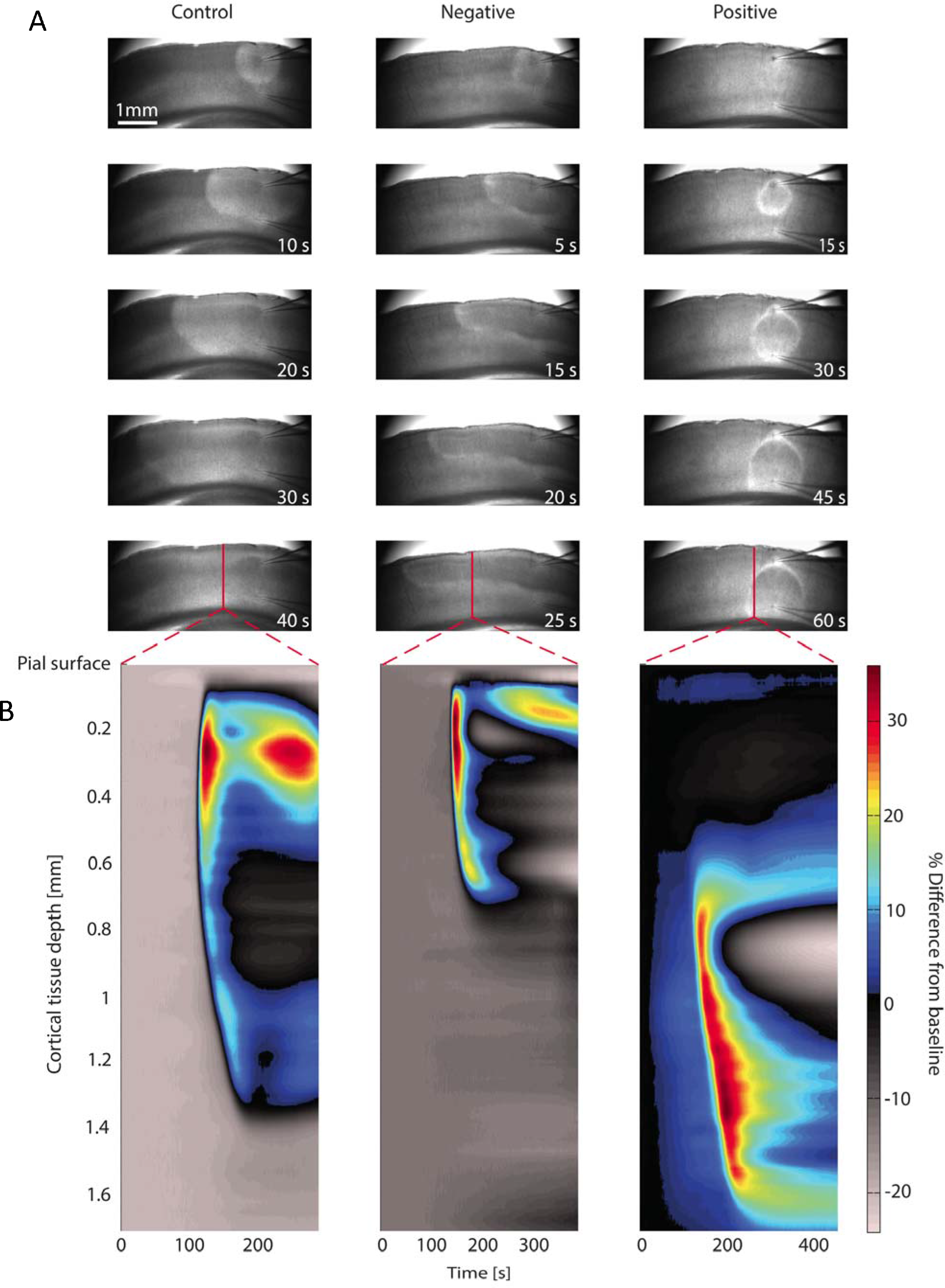
Examples of modulation experiments. (**A**) Electric field effects (116 mV/mm) on the propagation and invasion of SD into the various layers of coronal slices imaged via IOS. Normal SD propagation through all cortical layers during a control trial (left). SD propagation under applied surface-negative DC field confines the spread to the superficial cortical layers (middle), while SD propagation was forced into the deeper layers of cortex and arrested by applied surface-positive DC field (right). (**B**) Corresponding timing of SD signal as a function of cortical depth and time, the color bar representing light intensity changes as percent of baseline for the line of pixels indicated in red in upper plots (lower row in (**A**)). Note that the pixel cross-section for the positive field trial was taken nearer to the injection area due to the spatial arrest of SD.

A spatiotemporal representation of the IOS SD signal in Figure 2B illustrates different evolutions of the signal as a function of trial type and cortical depth. The leading edge of the IOS SD wavefront is located in the superficial layers during control trials in the left plot. The surface-negative field restricts the spatial depth of the IOS signal to the superficial layers blocking propagation into the deeper layers in the middle plot. In the right plot, the positive field demonstrates forcing of SD to the cortical depths where the SD IOS wavefront is more gradual in rise and prolonged in duration as it dissipates and propagation ceases.

Analysis of the SD propagation velocities in Figures 3A-3B, indicated that the presence of a positive field slows while a negative field increases propagation velocity. Factorial ANOVA confirmed that field polarity as well as slice age accounted for significant variance in the data (p<0.001, n=104 slices) with no significant interaction between factors (see Table S1). With significant effects on SD observed, experiments testing dose responses on field strength amplitude, as well as the effect of various field application delays from the start of SD were explored next to determine the extent of the modulation.

Dose response studies (n=32 slices for each dose condition) conducted on surface-positive field strength and stimulus delay time confirmed that as field strength increases and delay time shortens, the effective SD arrest rate increased and total SD affected areas decreased (Figure 3C-3D). For surface-positive applied fields the SD arrest rate was proportional to the field strength and field delay time (Figure 3D). Note also in Figure 3C the bimodal nature of the distributions at lower field strengths and longer delay times; indeed, SD propagations that were not arrested (gray dots) consumed more tissue on average than effective polarization trials at the same field strengths that arrested SD (black dots). As the field strength increases from weak (54 mV/mm) to strong (216 mV/mm), the number of SD events arrested (out of 32 SD trials for each tested condition) rose to 100% for field strengths greater than 160 mV/mm (Figure 3C). The dose response curves and the rates of SD propagation arrest are shown to be consistent across a range of field strengths and delay times in Figure 3D. We also found that prepolarization of the brain tissue in advance (15 seconds) of the high K+ injection could prevent the induction of SD (Figure 3C, right) in a field strength intensity-dependent manner (Figure 3D, right).

**Fig. 3.**
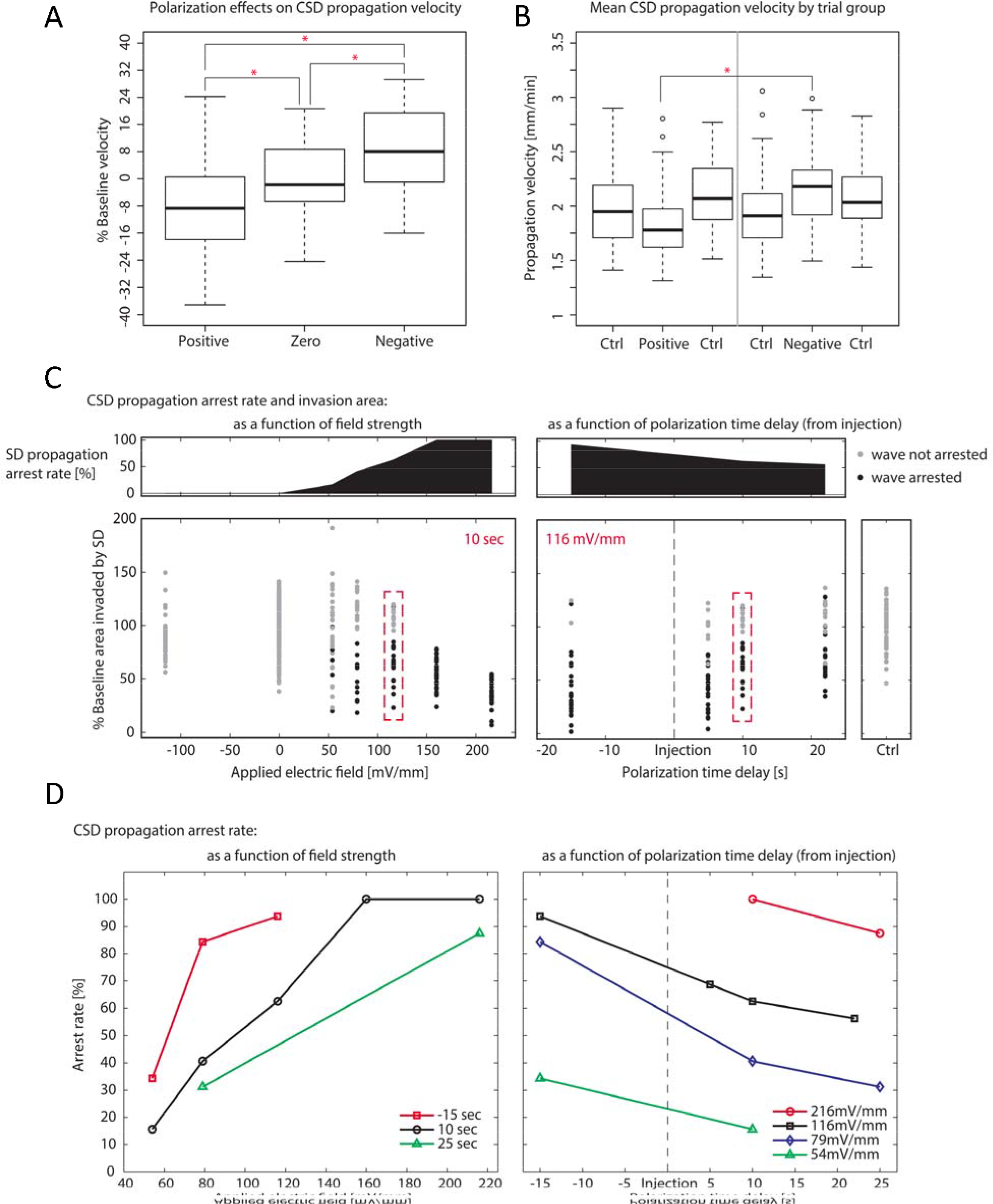
Summary of modulation experiments. (**A**) Summary of SD propagation velocity differences as a percentage of the averaged control trials for ±116mV/mm electric field strengths as a function of field polarity. The significant differences (p<0.01, n=104 slices) between groups are marked by asterisks. (**B**) SD propagation velocity as a function of each type of 3-epoch experiment (control with either positive or negative fields) for ±116mV/mm electric field strength. The flanking control trials for each polarization are shown here separately, while the gray dividing line indicates the independence between positive and negative experimental trials. The significant differences (p<0.01, n=104 slices) between groups are marked by asterisks, outliers are open circles. (**C**) Dose response of field strength efficacy (left), and stimulus timing from initiation of injection (right) in arresting SD propagation (arrest rate plotted above as the percentage of arrests out of 32 slices for each tested condition). Each distribution plotted for a different field strength and a different stimulus timing (for the 116 mV/mm field), compares the area of cortical tissue invaded by SD expressed as a percentage of the control trial SD invasion area. Black dots in the distribution indicate trials where the horizontal SD propagation was arrested by the applied field, gray dots indicate trials where the wave did not stop. Stronger fields and earlier applied fields are more effective in minimizing the tissue area affected by SD. Control trial plotted separately for comparison (lower far right). (**D**) Dose response of field strength (left) and stimulus delay time (right) efficacy in arresting SD propagation (as the percentage of arrests out of 32 slices for each commanded field or polarization time delay). Curves plotted for different stimulation delay times with respect to high K+ injection at time t=0. The time delay of the applied field response on the SD arrest rate is consistent across field strengths – the earlier the field is applied to the SD wave the more effective the arrest rate. Preconditioning the tissue with an applied field before SD induction shows the most effective arrest rates for each field strength tested. The effectiveness of stopping SD was proportional to the strength of the applied field at all delay times.

#### High K+ bath perfusion ignition of SD

Repetitive periodic SD can be produced in very high concentrations of of extracellular K+, from 26 mM (*29*) to 40 mM (*30*). In high K+ perfusion experiments, applied negative fields showed a dramatic near-simultaneous ignition of SD along the superficial layers in front of the wavefront, whereas positive fields prevented further SD invasion into the superficial half of the cortex for a about 30 seconds (Figure S1). Within 30 seconds the high K+ bath depolarization of those tissues seemed to overwhelm the temporary blockage and SD collapsed inward on the unaffected region and consumed the rest of the tissue (frames 5 and 6 in right column of Figure S1). Also notable in the lower right side of the fourth frame in Figure S1 is a second SD wave ignition in deeper layers of the cortex caused by depolarization of the deeply located neuronal membranes by surface positive polarization in the presence of high bath K+. We note that SD can be reliably produced in tangential cortical slices, which have the advantage of a more isotropic preparation of cortex (Figure S2). Our collective experience with high K+ induced SD in coronal (n=77) and tangential (n=10) slices suggested that the effects of DC polarization of the cortex was counteracted by the effects of excessively high K+ bath perfusion on the cortical tissue therefore our main results were studied with local high K+ injections to ignite SD. In brains of animal or human patients experiencing SD, the extracellular space is not known to have such pathologically high concentrations of K+ unless an SD wave has been triggered. Indeed, there appears a physiological ceiling for extracellular K+ between 12-15 mM that is rarely exceeded except during SD (*31*).

### Discussion

In this work, we report experimental findings consistent with biophysical computational models (*24*) – that polarization of the neuron along the somatodendritic axis could be utilized to speed up, slow down, arrest and even prevent propagating waves of SD. Consistent with the biophysical modeling (*24*), but nonetheless surprising, is that the polarity that blocks SD is opposite to that which suppresses spikes (*3, 4, 15*) and seizures (*11*).

Our IOS and K+ optical measurements suggests multiple effects on SD. First, intracellular hyperpolarization of the apical dendrites could counteract SD triggering that depends upon depolarization of the dendrites (*24, 25*). Second, surface-positive polarization might act in an electrophoretic manner to drive extracellular positive charges including K+ away from superficial dendritic membranes particularly sensitive to depolarization by extracellular K+ during SD ignition. Third, surface-positive fields may contribute to the increased spatial buffering of local elevated K+ by glia. These effects might act synergistically to block further propagation of SD, and are consistent with the suspected role of superficial cortical layer dendritic membranes in the horizontal spread of SD (*32, 33*). It is suspected that the spatial extent of dendritic arborizations and high surface area to volume ratio of dendritic membranes in these superficial layers, combined with the relatively small extracellular volume fraction, contributes to their sensitivity to high concentrations of extracellular K+ during the ignition and propagation of SD (*34*). It is felt that this sensitivity explains the preference of SD to propagate along these layers as shown in Figures 2 and S1.

From past polarization experiments during spiking (*4, 17*) and seizure (*11, 12, 19*) activity, one can generate an increase in such activity with surface-positive fields as applied herein which depolarize cortical somata (the opposite is seen with the inverted geometry of the hippocampal principal cells, see (*15*)). Yet such experiments in normal cortex and with seizures failed to generate consistent SD. In our present experiments, despite mildly elevated K+ in the perfusate (6.25 mM) that helped support propagating SD, our data shows no overt evidence that seizures were generated by the applied positive fields. Control trials following positive field applications demonstrated full tissue recovery and readily generated SD, whereas SD does not invade regions with active or recent seizure activity (*35*). Neurons involved with the SD wavefront were likely inactivated by the local elevation in extracellular K+ causing depolarization block (*36*). The observation that applied positive fields to arrest SD might not cause seizures under conditions where SD develops, and that negative fields do not cause SD under conditions where seizures are generated (*11–13*), raises the intriguing prospect that the control required to modulate either condition (SD or seizures) might be state-dependent. Indeed, recent work has demonstrated that there is a unification between spikes, seizures, and SD that are all part of the dynamical repertoire of the neuronal membrane, but that such activities are each confined to separate regions of the parameter space (ion concentrations, oxygenation, cell volume status) that defines which normal or pathological state might arise (*23, 31*). Our present results suggest that control of seizures or SD might require that the state of the brain tissue be estimated, so that the appropriate polarity and type of stimulation can be applied.

### Materials and Methods

#### Experimental design

We performed a series of experiments triggering SD with local high K+ injection (*37, 38*). This local injection of K+ supports SD propagation in perfusates with much lower concentrations of K+ (6.25 mM) than high K+ perfusion methods, and permits controlled single SD wave generation (*29*). The experimental protocol was built from 3-epoch trials as shown in Figure 1C. First a control trial was performed with no field present to assess the health and condition of the slice – any slice that appeared to be in poor condition optically or unable to propagate a strong SD signal was discarded. Slices with a strong first SD signal were allowed to recover for 15 minutes, then a second polarization trial was conducted with the electric field on, and following a second recovery period a third SD control trial was performed to control for temporal effects and isolate polarization effects related to the field. Repetitive SD episodes for any given slice were limited to these 3-epoch paradigms (one experimental trial between two controls) to ensure slice viability and maximize the number of healthy slices utilized per experiment. All experiments were performed in accordance with and approval from the institutional animal care and use committee of The Pennsylvania State University. Neocortical slices were obtained from male Sprague-Dawley rats aged P16 to P21. Briefly, the animals were deeply anesthetized with diethyl-ether and decapitated, the brain removed and coronal slices were sectioned from occipital neocortex. The slices were prepared differently for each of the experiments and are described in the following sections.

#### Slice preparation

After decapitation, the whole brain was quickly and carefully removed to chilled (4°C) artificial cerebrospinal fluid (ACSF) for 60 seconds containing the following (in mM): 126 NaCl, 2.5 KCl, 2.4 CaCl2, 10 MgCl2, 1.2 NaH2PO4, 18 NaHCO3, and 10 dextrose. ACSF was saturated with 95% O2 and 5% CO2 at room temperature for 1 hour before the dissection with an osmolality ranging from 295-310 mOsm and pH from 7.20-7.40. Coronal occipital cortical slices were cut with a vibratome on the rostro-caudal and mediolateral coordinates of bregma −2 to −8 mm and lateral 1 to 6 mm, respectively. The first cut was made 1100 μm deep from the caudal surface and discarded, for bath applied high K+ experiments 3 slices were taken from each hemisphere each 300 μm thick while for locally injected high K+ experiments, 4 slices were taken from each hemisphere each 450 μm thick. Tangential slices were sectioned with the first cut made 100 μm deep from the pial surface and discarded, and a 500 μm thick slice was taken of the middle cortical layers (*39*). After cutting, slices were transferred to a chamber containing ACSF and recovered for 30 minutes at 32-34°C then incubated at room temperature (20-22°C) for an additional 30 minutes prior to recording. For potassium dye loading, Asante Potassium Green-2 AM ester (Abcam) was dissolved in DMSO and 0.045% Pluronic F-127 (TEF labs) and added to the incubation ACSF for a final concentration of 20 μM, followed by 1 hour of loading and 1 hour of rinsing at room temperature before imaging.

#### SD generation

A schematic of the recording chamber and arrangement of electrodes is shown in Figure 1. For local injection induction of SD, modified ACSF (in mM) 121.25 NaCl, 6.25 KCl, 1.5 CaCl2, 0.5 MgCl2, 1.25 NaH2PO4, 25 NaHCO3, and 25 dextrose was perfused over the slice at a rate of 2.8-3.0 mL/min at 30°C during recording. To evoke SD, a pico-liter injector (PLI-10; Warner Instruments) was used to inject 3 M KCl into layer II/III using a 150 ms pulse at a pressure of 28.4 psi (which was more than sufficient to ignite robust spreading depression propagation).

For bath applied high K+ induction, 26 mM KCl replaced equimolar NaCl and MgCl2 was lowered to 1.3 mM. The flow of ACSF into the chamber was switched to 26 mM K+ solution; SD would typically initiate from one or more foci in layers II/III of the cortex after 1-2 minutes of exposure at which point the flow was switched back to normal ACSF. After each SD episode, the slice was allowed to recover for 15 minutes and in this way multiple SD waves could be repeatedly evoked using either method without apparent damage to the tissue.

#### Electric field application

The recording chamber (RC-22C; Warner Instruments) was fitted with Ag/AgCl sintered pellet electrodes (4.37 mm x L, 1.20 mm x H, separation distance 5.60 mm) insulated such that only the tissue in the camera’s field of view would be polarized as shown in Figure 1A. Extracellular pipettes (4-6 MΩ) were pulled from thick-walled borosilicate glass capillaries with filaments (OD=1.5 mm, ID=0.86 mm; Sutter Instruments) and filled with 3 M KCl. The injection pipette was inserted into the middle top of layer II/III with the second pipette located directly inline ∼900 μm in the deeper layers of the tissue near the bottom of the cortex. Differential voltage measurements were acquired (MultiClamp 700A; Axon Instruments) as a feedback measurement of the field inside the tissue. In order to guarantee the field strength a Labview interface was used to control a National Instruments BNC-2120 signal generator and drive voltage commands to a constant current isolated stimulator (Model 2200; AM-Systems) as in Figure 1B. A proportional integral differential (PID) feedback control system was implemented on the differential voltage measurement measured within the cortical tissue, which allowed precise and reproducible electric field strengths to be commanded for each stimulus trial (Figure 1C).

#### Spreading depression imaging via the intrinsic optical signal (IOS) and APG-2 epi-fluorescence

SD was observed by measuring the changes in light transmittance through the tissue, also known as the intrinsic optical signal (IOS), which is related to local neuronal activity and cell swelling. In our configuration, the transmitted light of our IOS is dominated by cell swelling during SD (*29*). Optical imaging and local field potential were recorded simultaneously during a portion of the experiments; slices were trans-illuminated with white light filtered at 757 ± 30 nm. For simultaneous IOS and epi-fluorescence imaging, a Dual-View optical multichannel system (Photometrics, Tucson, AZ) was configured as in Figure 1D. Video frames were acquired with a 4x objective at 30 Hz (0.5 Hz for simultaneous IOS/epi-fluorescence, with a 300 ms illumination) using a 1296 × 966 pixel charge-coupled device (CCD) camera with 12-bit resolution (acA1300-30um; Basler) which was set for 1x camera amplifier gain and stored on disk for off-line analysis. Each pixel in the images equated to a 4 × 4 μm square, the full resolution providing a 5.184 × 3.864 mm imaging window of the cortical tissue.

#### Data analysis

Video data were spatially filtered with a Gaussian smoother (10×10 pixel kernel) then filtered forward and backward in time with a 3rd order Butterworth filter at 0.25 Hz and analyzed in MATLAB (Mathworks, Natick, MA). The filtered time series of luminous intensity at each pixel was then analyzed for the SD wavefront by taking the temporal derivative and finding the time point of maximum rate of change. This ensemble of SD wavefront detection time points at each pixel creates a map of where the SD front was detected and during what time point it passed each pixel, in short a “time map” containing spatiotemporal information about the propagation of the IOS (or APG-2 dye) signals of spreading depression. From this measurement, SD propagation velocity and the extent of invasion into the tissue could be computed and further analyzed for the effects of various applied electric fields during SD propagation.

#### Statistical analysis

Factorial ANOVA was used to test for significant effects of polarity type and slice age on propagation velocity (see Table S1). Since successive SD events display a trend of increasing propagation velocity (as seen from the increase in average velocity from first to last control trials in Figure 3B), we normalized the velocity of each slice polarity trial as a percentage of the average of the corresponding controls. All polarity groups were tested for normality (Shapiro-Wilk test, W=0.9879, p=0.224, and graphically in Figures S3 and S4) and homogeneity of variance (Bartlett’s test, T=0.8512, df=2, p=0.6531) before being compared using Bonferroni corrected paired t-tests where p<0.01 was regarded as significant (asterisks in Figure 3A).

## Acknowledgments

**General**: We are grateful for the many helpful discussions with Koen Dijkstra, Stephan van Gils, and Stephen van Wert.

**Funding:** Supported by US-German Collaborative Research in Computational Neuroscience program: NIH Grant 1R01EB014641 (AW, BJG, SJS), and Bundesministerium für Bildung und Forschung, BMBF 01GQ1001B (MD), and NIH BRAIN Initiative 1R21EY026438 (HK, SJS).

**Author contributions:** AJW, YX and HK performed all brain slice experiments, BJG and SJS aided in the experimental design and implementation of the experimental data collection system, and MD assisted with the analysis of the video data.

**Competing interests:** The authors disclose no conflicts of interest.

**Data and materials availability:** Data relevant to the results presented in this paper are archived online at (placeholder).

## Supplementary Figures

**Fig. S1.**
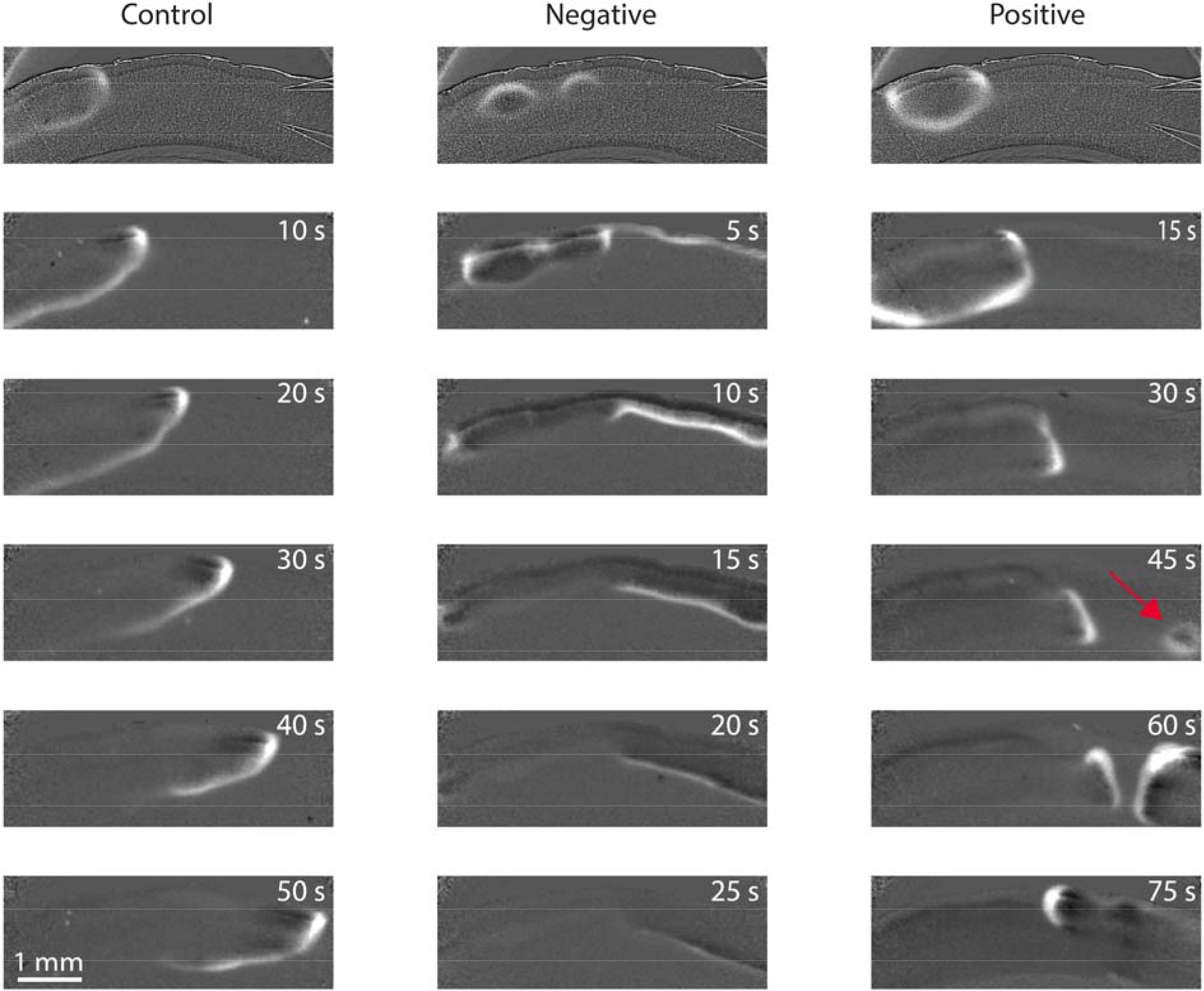
Electric field effects on the propagation and invasion of SD into the various layers of coronal slices evoked spontaneously in high K+ bath perfusion (26 mM), imaged via IOS. Normal SD propagation through all cortical layers during a control trial (left). SD propagation under surface-negative DC field (center) applied just before the second frame causes SD ignition to rapidly occur simultaneously across the uppermost cortical layers and propagation continues unabated. SD propagation was temporarily blocked from invading upper layers of cortex by an applied positive DC field (right), which in high K+ bath caused a secondary SD wave ignition in the deeper layers of cortex (red arrow). Eventually the continuous cellular depolarization from high K+ bath causes SD invasion to overcome the blocking effects of the positive field. Images contrast enhanced for display by background intensity subtraction.

**Fig. S2.**
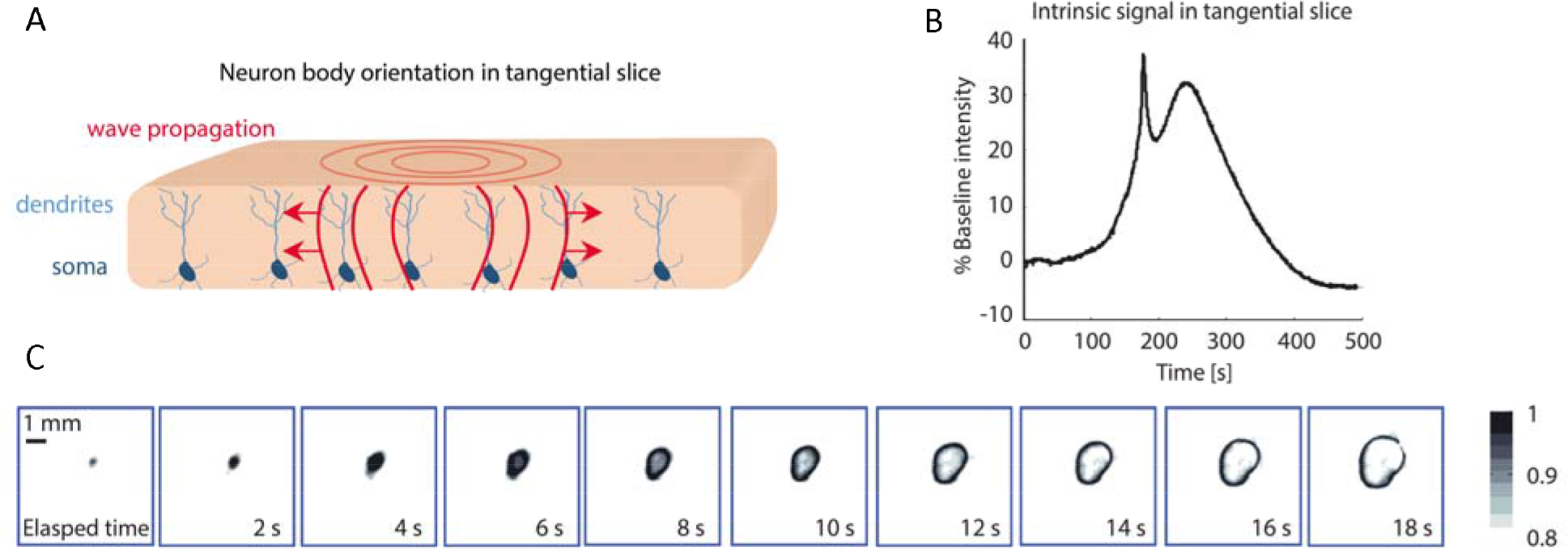
Summary of SD propagation in tangential slices. (**A**) Diagram of a tangential cortical slice and neuron body orientation demonstrating the typical propagation of SD and the near isotropic nature of tangential slices. (**B**) Intrinsic optical signal (IOS) recorded in tangential slices comprising a fast SD propagation related intensity peak followed by a slower recovery signal component. (**C**) IOS signal of ring wave propagation in tangential slices, darker color represents increasing depolarization.

**Fig. S3.**
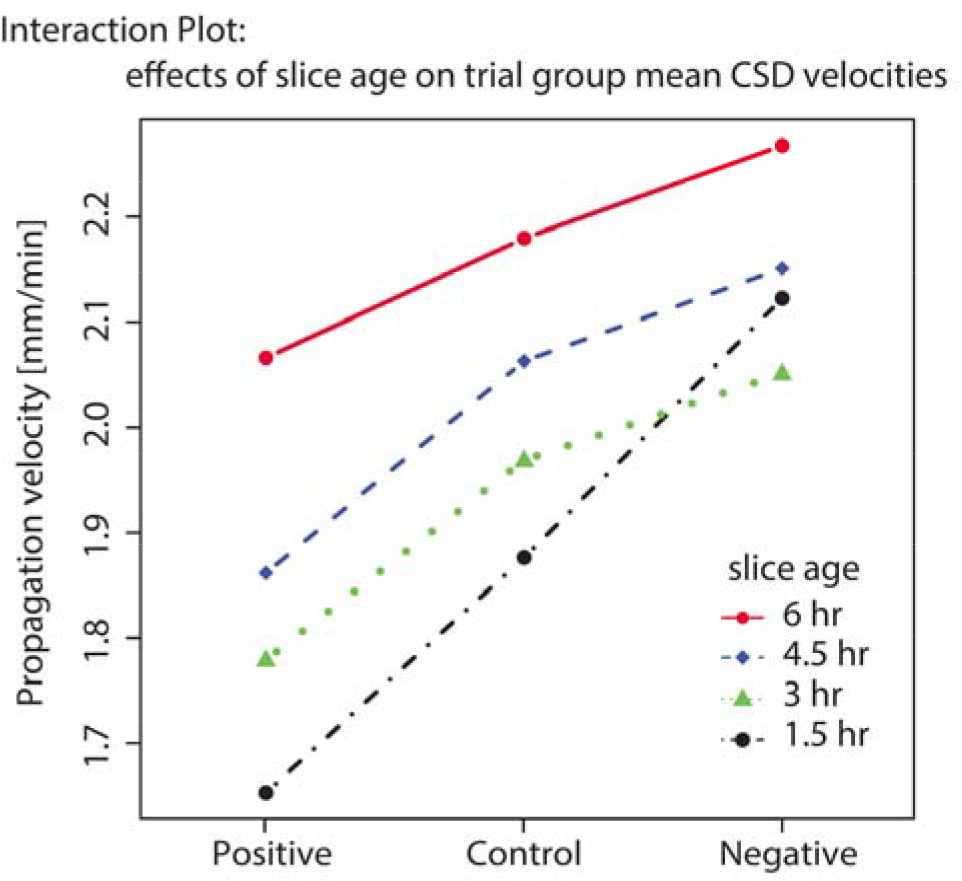
Slice aging effects on the average propagation velocity of SD where each point represents an average of 13 slices. Note the consistency of slice age and polarity data factors as the overall trend between polarity groups remains intact as slice age varies. Slice aging results in faster SD propagation.

**Fig. S4.**
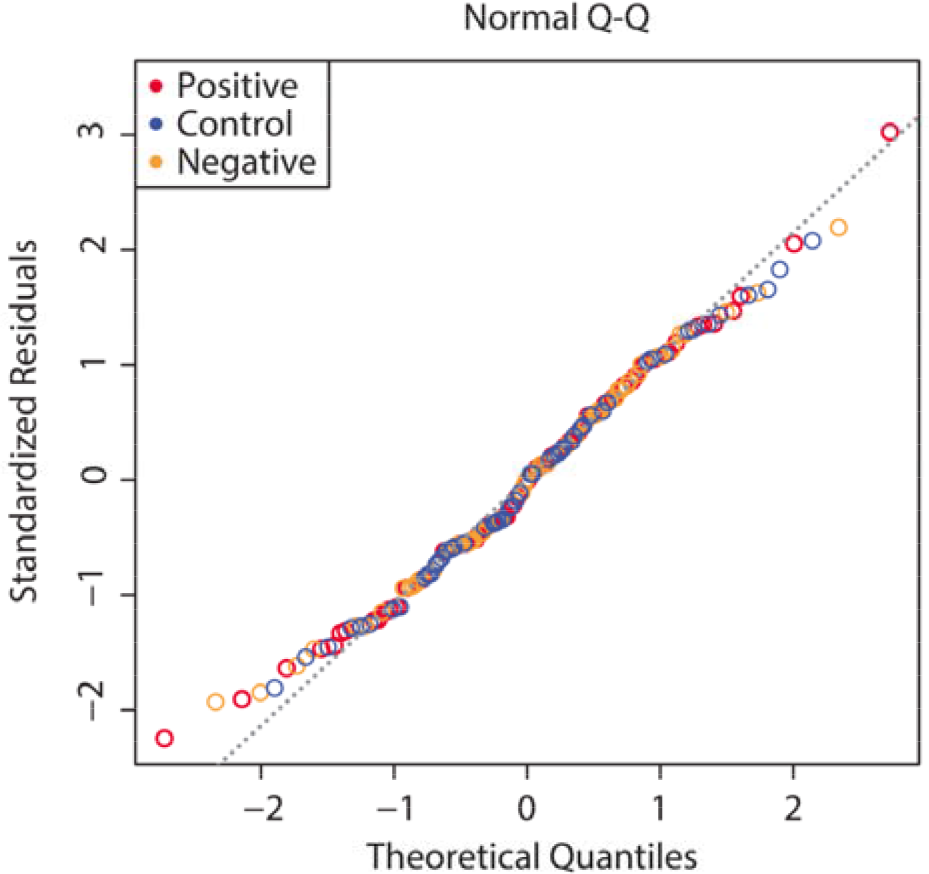
A normal quantile-quantile plot testing the normality of the propagation velocity data.

**Table S1.** Factorial ANOVA on Mean CSD Propagation Velocity.

|  | *Df* | *Sum Sq* | *Mean Sq* | *F value* | *P Value* | *Sig.* |
| --- | --- | --- | --- | --- | --- | --- |
| *Slice age* | 3 | 3.839 | 1.2797 | 13.503 | 2.87e-08 | *** |
| *Polarity* | 5 | 3.545 | 0.7091 | 7.483 | 1.28e-06 | *** |
| *Slice age/Polarity* | 15 | 0.742 | 0.0495 | 0.522 | 0.928 |  |
| *Residuals* | 288 | 27.293 | 0.0948 |  |  |  |

*Significance: p<0.001\*\*\**

## References and Notes

1. W. A. Rushton, The effect upon the threshold for nervous excitation of the length of nerve exposed, and the angle between current and nerve. J. Physiol. 63, 357–77 (1927).

2. C. A. Terzuolo, T. H. Bullock, Measurement of imposed voltage gradient adequate to modulate neuronal firing. Proc. Natl. Acad. Sci. U. S. A. 42, 687–94 (1956).

3. D. P. Purpura, J. G. McMurtry, Intracellular activities and evoked potential changes during polarization of motor cortex. J. Neurophysiol. 28, 166–185 (1965).

4. J. G. Jefferys, Influence of electric fields on the excitability of granule cells in guinea-pig hippocampal slices. J. Physiol. 319, 143–152 (1981).

5. A. R. Gardner-Medwin, A study of the mechanisms by which potassium moves through brain tissue in the rat. J. Physiol. 335, 353–374 (1983).

6. C. Y. Chan, C. Nicholson, Modulation by applied electric fields of Purkinje and stellate cell activity in the isolated turtle cerebellum. J. Physiol. 371, 89–114 (1986).

7. C. Y. Chan, J. Hounsgaard, C. Nicholson, Effects of electric fields on transmembrane potential and excitability of turtle cerebellar Purkinje cells in vitro. J. Physiol. 402, 751–71 (1988).

8. D. M. Durand, M. Bikson, Suppression and control of epileptiform activity by electrical stimulation: a review. Proc. IEEE. 89, 1065–1082 (2001).

9. P. Krack et al., Five-Year Follow-up of Bilateral Stimulation of the Subthalamic Nucleus in Advanced Parkinson’s Disease. N. Engl. J. Med. 349, 1925–1934 (2003).

10. T. Wichmann, M. R. DeLong, Deep Brain Stimulation for Neurologic and Neuropsychiatric Disorders. Neuron. 52, 197–204 (2006).

11. B. J. Gluckman et al., Electric field suppression of epileptiform activity in hippocampal slices. J. Neurophysiol. 76, 4202–5 (1996).

12. B. J. Gluckman, H. Nguyen, S. L. Weinstein, S. J. Schiff, Adaptive electric field control of epileptic seizures. J. Neurosci. 21, 590–600 (2001).

13. M. Bikson et al., Suppression of epileptiform activity by high frequency sinusoidal fields in rat hippocampal slices. J. Physiol. 531, 181–191 (2001).

14. D. Tranchina, C. Nicholson, A model for the polarization of neurons by extrinsically applied electric fields. Biophys. J. 50, 1139–56 (1986).

15. D. P. Purpura, A. Malliani, Spike generation and propagation initiated in dendrites by transhippocampal polarization. Brain Res. 1, 403–6 (1966).

16. J. G. Jefferys, Nonsynaptic modulation of neuronal activity in the brain: electric currents and extracellular ions. Physiol. Rev. 75, 689–723 (1995).

17. J. T. Francis, B. J. Gluckman, S. J. Schiff, Sensitivity of neurons to weak electric fields. J. Neurosci. 23, 7255–61 (2003).

18. K. A. Richardson, S. J. Schiff, B. J. Gluckman, Control of traveling waves in the Mammalian cortex. Phys. Rev. Lett. 94, 28103 (2005).

19. K. A. Richardson et al., In Vivo Modulation of Hippocampal Epileptiform Activity with Radial Electric Fields. Epilepsia. 44, 768–777 (2003).

20. A. Leao, Spreading depression of activity in the cerebral cortex. J Neurophysiol. 7, 359–390 (1944).

21. M. Lauritzen, Pathophysiology of the migraine aura. The spreading depression theory. Brain. 117 ( Pt 1, 199–210 (1994).

22. M. Lauritzen et al., Clinical relevance of cortical spreading depression in neurological disorders: migraine, malignant stroke, subarachnoid and intracranial hemorrhage, and traumatic brain injury. J. Cereb. Blood Flow Metab. 31, 17–35 (2011).

23. Y. Wei, G. Ullah, S. J. Schiff, Unification of Neuronal Spikes, Seizures, and Spreading Depression. J. Neurosci. 34, 11733–11743 (2014).

24. H. Kager, W. J. Wadman, G. G. Somjen, Conditions for the triggering of spreading depression studied with computer simulations. J. Neurophysiol. 88, 2700–12 (2002).

25. J. Makarova, M. Gómez-Galán, O. Herreras, Variations in tissue resistivity and in the extension of activated neuron domains shape the voltage signal during spreading depression in the CA1 in vivo. Eur. J. Neurosci. 27, 444–56 (2008).

26. B. Grafstein, Mechanism of spreading cortical depression. J. Neurophysiol. 19, 154–71 (1956).

27. V. I. Koroleva, J. Bures, Blockade of cortical spreading depression in electrically and chemically stimulated areas of cerebral cortex in rats. Electroencephalogr. Clin. Neurophysiol. 48, 1–15 (1980).

28. F. J. Brinley, E. R. Kandel, W. H. Marshall, Potassium outflux from rabbit cortex during spreading depression. J. Neurophysiol. 23, 246–256 (1960).

29. T. R. Anderson, R. D. Andrew, Spreading depression: imaging and blockade in the rat neocortical brain slice. J. Neurophysiol. 88, 2713–25 (2002).

30. N. Zhou, G. R. J. Gordon, D. Feighan, B. A. MacVicar, Transient Swelling, Acidification, and Mitochondrial Depolarization Occurs in Neurons but not Astrocytes during Spreading Depression. Cereb. Cortex. 20, 2614–2624 (2010).

31. G. Ullah, Y. Wei, M. A. Dahlem, M. Wechselberger, S. J. Schiff, The Role of Cell Volume in the Dynamics of Seizure, Spreading Depression, and Anoxic Depolarization. PLOS Comput. Biol. 11, e1004414 (2015).

32. B. Grafstein, Locus of propagation of spreading cortical depression. J. Neurophysiol. 19, 308–16 (1956).

33. S. Ochs, K. Hunt, Apical dendrites and propagation of spreading depression in cerebral cortex. J. Neurophysiol. 23, 432–44 (1960).

34. B. de Luca, J. Bures, Development of cortical spreading depression and of its transition to the caudate nucleus in rats. Dev. Psychobiol. 10, 289–97 (1977).

35. V. I. Koroleva, J. Bures, Circulation of cortical spreading depression around electrically stimulated areas and epileptic foci in the neocortex of rats. Brain Res. 173, 209–15 (1979).

36. M. M. Reddy, J. Bureš, Cortical (K+)e and the stimulation induced blockade of spreading depression in the rat cerebral cortex. Neurosci. Lett. 17, 243–247 (1980).

37. S. Canals et al., Longitudinal depolarization gradients along the somatodendritic axis of CA1 pyramidal cells: a novel feature of spreading depression. J. Neurophysiol. 94, 943–51 (2005).

38. A. Tottene et al., Enhanced excitatory transmission at cortical synapses as the basis for facilitated spreading depression in Ca(v)2.1 knockin migraine mice. Neuron. 61, 762–73 (2009).

39. X. Huang et al., Spiral waves in disinhibited mammalian neocortex. J. Neurosci. 24, 9897–902 (2004).

